# Enhancing the versatility of photocrosslinkable silk fibroin using an eco-friendly solvent

**DOI:** 10.1101/2024.10.06.616881

**Authors:** Anne Katherine Brooks, Vamsi K. Yadavalli

## Abstract

Silk fibroin (SF), known for its biocompatibility and versatility, has been widely studied in tissue engineering and biomedical devices. The modification of silk fibroin with photoreactive groups has been used to create novel biomaterials that undergo a liquid-to-solid transition upon exposure to light, enabling precise control over structure formation, pore geometry, and degradation. This advancement of photofibroin (PF) has been shown for the biofabrication of hydrogels, 3D scaffolds, and micro-patterned surfaces suitable for biomedical applications, including tissue scaffolds and bioelectronics. Here, we present a further improvement using a water based ternary solvent of calcium chloride-ethanol-water (Ajisawa reagent (AR)), to dissolve photofibroin, offering a sustainable alternative to previously used organic solvents. PF in AR is shown to be compatible with various light-based manufacturing techniques including soft lithography, photolithography, and 3D printing, enabling the fabrication of multiscale structures with high fidelity. The gels formed demonstrate excellent cytocompatibility, supporting cell adhesion and growth without additional coatings, making them ideal for regenerative medicine. The integration of conductive polymers, such as PEDOT:PSS as a 3D printable conducting gel opens possibilities for bioelectronics. The research represents a significant step forward in employing the versatile photofibroin as a sustainable, high-performance biomaterial for diverse applications.

## 1. Introduction

Naturally derived materials including collagen, gelatin, silk proteins (e.g. fibroin and sericin), alginate, and chitosan have been widely studied and used for diverse applications due to attractive properties including sustainable sourcing, biorecognition, biocompatibility, and often, biodegradability.[1-3] Silk fibroin (SF) is a particularly versatile (bio)material owing to high mechanical strength, processability, controlled biodegradation, and ability to be functionalized.[4-6] Applications in tissue engineering scaffolds, drug delivery, and wound healing have been reported over the past few decades.[7, 8] Recently, durable, mechanically flexible and lightweight micro-scale devices and components have been shown.[9, 10] With techniques such as soft lithography, micro-molding, and electrospinning, SF has been used to form cell scaffolds as well as microdevices and biosensors.[11] Concurrently, light-based fabrication has become increasingly popular for constructing micro- and nanoscale structures.[12] Light (often ultraviolet (UV)) can enable fine control over feature sizes and shapes to form complex multi-layered devices with optimized performance and compatibility with biological systems. Photolithography, stereolithography (SLA) and digital light printing (DLP) are among highly scalable and high-throughput light-based “writing” techniques for the creation of precise and intricate patterns in 2D and 3D.[13, 14] However, such tools require materials with the ability to undergo a phase transition from solid to liquid (or vice-versa). SF has been physically crosslinked using water annealing, or immersion in methanol or ethanol to induce changes in protein structure. However, such physical crosslinking results in limited form factors (e.g. hydrogels) and unpredictable gelation (temporal and spatial).[15] Chemical crosslinking agents (e.g., diglycidyl ether epichlorohydrin, ethylene glycol, glutaraldehyde, genipin etc. [16]) have limited temporal control, or potentially introduce cytotoxicity. Alternatives to both physical and chemical crosslinking are desirable. Photocrosslinkable materials were developed to provide controlled polymerization/de-polymerization using light, thereby enhancing safety profile as well as fabrication versatility.

Photocrosslinkable silk fibroin (PSF, or photofibroin (PF)) refers to modification of natural SF with photo-reactive groups providing the light-induced phase transition. In this work, we focus on the liquid to solid transition (crosslinking) resulting in a negative tone photoresist-like material. Photocrosslinking allows precise control over the material’s properties including pore geometry and degradation, while enabling the fabrication of complex and well-defined structures such as hydrogels, light-based additive manufacturing for 3D scaffolds, and micro-patterned surfaces. The material is crosslinked upon exposure to specific wavelengths of light – typically, UV (365 nm) or blue (405 nm). PF can be used with photolithography or with techniques such as etching and deposition, to produce complex biodevices that combine the inherent biocompatibility and biodegradability of silk with the precision and versatility of light-actuated techniques.

Our group was one of the first to report on photocrosslinkable forms of both fibroin and sericin (PF and PS), using chemical modification of the amino acid side chains via 2-isocyanatoethyl methacrylate (IEM) (**Figure 1**).[18, 19] Since our report, syntheses of other variants of photocrosslinkable silk fibroin have been reported using reagents such as methacrylic anhydride (MA) [20], glycidyl methacrylate (GMA),[21] and carbic anhydride.[22] An excellent review of these materials recently outlined the various chemical functionalization strategies and characterization for photocrosslinkable SF.[23] The corresponding materials have been variously referred to as methacrylated silk fibroin (Sil-MA), silk fibroin methacrylate (Sil-IEM), norbornene-functionalized silk fibroin (SF-NB) or silk fibroin methacrylate (Sil-GMA). Of these, the Sil-MA form is perhaps the most used. Sil-MA forms hydrogels with covalently bonded networks through photoinitiated radical polymerization by UV light in the presence of a photoinitiator. An advantage of Sil-MA is the polymerization which can be performed under moderate conditions (room temperature, neutral pH, and in aqueous environments). Together with its cytocompatibility and adaptability to 3D printing, it has also recently been commercialized.

**Figure 1.**
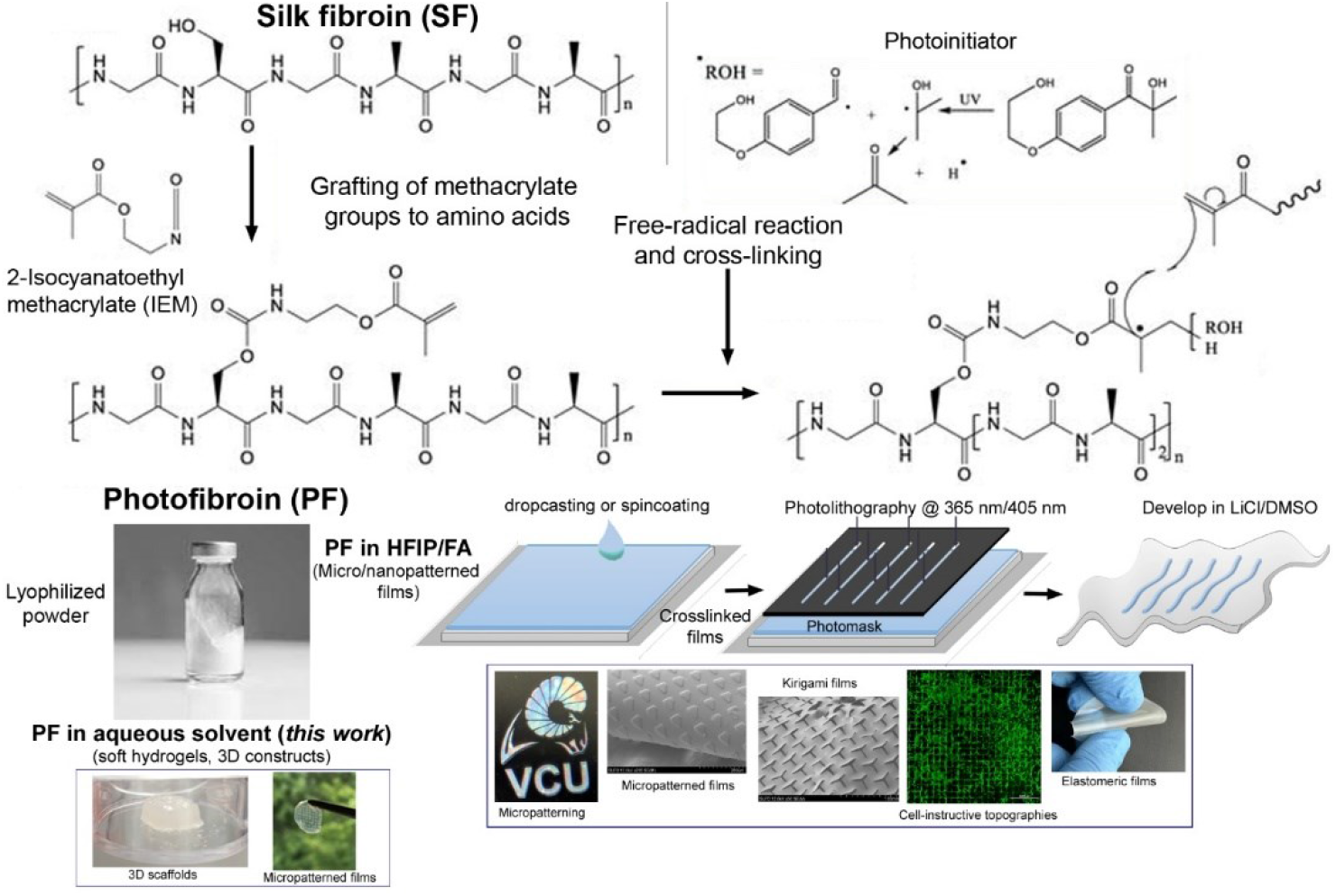
Synthesis of the photofibroin via IEM grafting and its crosslinking in the presence of a photoinitiator. Adapted from [17, 18]. Two different routes for utilization of the PF biomaterial are shown below. The formation of high precision photolithography-based forms is enabled by dissolution in HFIP/FA. In this work, the use to form softer hydrogels via an aqueous solvent is shown.

While Sil-MA is widely used, it tends to have a lower mechanical strength, leading to various reports using other materials as composite bioinks.[23] While some groups have had success printing large and multilayered constructs,[24] others have not – primarily due to loss of resolution or collapse of structures.[25, 26] Combinations with other (bio)polymers have often been sought to remedy this.[27] Due to the expanded capability of reactive IEM, higher degrees of methacrylation are obtained than in the case of MA, with values up to 9.3% reported. Thus, the photofibroin (PF) formed via IEM can form structures of extraordinary fidelity and strength. PF use has been limited to forming solutions using (hexafluoroisopropanol) HFIP or formic acid (FA) (**Figure 1**) which have been excellent at photolithographic routes of fabrication, with the solvents not contributing negatively to the bio/cytocompatibility of the constructs. However, the use of HFIP/FA can also be a deterrent to wider use. Here, we show an enhancement of the profile of PF, by demonstrating Ajisawa’s reagent (AR) as a water-based solvent.[28] AR was developed nearly 30 years ago and is a ternary solvent system consisting of calcium chloride, DI water, ethanol (EtOH) dissolved in a 1:8:2 molar ratio. The photopolymer solution can be micromolded into large volume and high-fidelity structures via soft lithography, or rapidly 3D printed using digital light processing (DLP). We show that the use of the ternary solvent preserves the cytocompatibility of photofibroin while conferring added adhesiveness under hydrated conditions. For the first time, we are also able to successfully demonstrate a 3D printed conducting grid comprising photofibroin and the conducting polymer PEDOT:PSS in an aqueous bath. The combination of these versatile enhancements makes the ternary solvent-photofibroin system optimal as a robust biomaterial for diverse applications including cell growth and tissue regeneration.

## 2. Materials and Methods

### 2.1. Synthesis of photofibroin and photoreactive precursor solution

Photofibroin (PF) was prepared as described previously.[29] Silk fibroin fibers were extracted from *Bombyx mori* cocoons via degumming, and regenerated to obtain fibroin powder.[30] Photoreactive methacrylate groups were conjugated to the fibroin protein via isocyanate addition using 2-isocyanatoethyl methacrylate (IEM) to form photofibroin (PF).[18] PF is not water soluble but was previously shown to be soluble in hexafluoroisopropanol (HFIP) and formic acid (FA) to form a precursor solution. In this work, Ajisawa’s reagent (AR) is used as a water-based solvent for PF,[28] with calcium chloride (Sigma Aldrich, anhydrous), DI water, ethanol (EtOH) dissolved in a 1:8:2 molar ratio. PF can be fully dissolved in AR at over 10 % w/v by breaking apart PF granules into a fine powder, adding reagent and vortexing.

Since CaCl_2_ is hygroscopic, piece-wise addition of a fine powder together with vortexing enhances dissolution without clumping. If it is desired to reduce the amount of CaCl_2_ used in solution, the PF solubility decreased below 3:32:8 of CaCl_2_:water:EtOH molar ratio (0.75:8:2). This (lower bound) ratio, denoted as AKR, was used interchangeably with AR in this study to form photoreactive precursor solutions. Different photoinitiators were used to enable crosslinking at various wavelengths. Irgacure 2959 is easily soluble in AR/AKR solution and enables crosslinking at 365 nm. For crosslinking at 405 nm, lithium phenyl-2,4,6-trimethylbenzoylphosphinate (LAP) is used. However, LAP is not soluble in either AR/AKR. The desired amount of LAP (0.5% w/v) is solubilized in a small volume of DMSO (<50µL) and then dissolved in the PF solution to form a photoreactive precursor solution. To form a conductive hydrogel precursor, the conducting polymer poly(3,4-ethylenedioxythiophene)-poly(styrene sulfonate) (PEDOT:PSS) was dispersed in the solvent before adding PF via ultrasonication to yield 0.25% or 0.5% w/v in the final solution.

### 2.2. Microfabrication of silk fibroin hydrogels

Hydrogel precursor solutions were cast into circular molds on glass slides (50 µL solution into a 0.8 cm diameter circular well) and cured in a Loctite Zeta 7401UV Chamber (Henkel Canada Corporation) for 3 minutes to yield crosslinked hydrogels. For patterning via soft lithography, solution is cast into PDMS micromolds coated in Sigmacote^®^ siliconizing reagent (Sigma Aldrich) for easy removal. Features included 500 µm conical spikes, 100 µm troughs, and macroscale shapes. Multi-layered patterning was possible by casting and crosslinking conductive solution in micropatterned grooves, crosslinking, then casting and crosslinking a second layer on top. Removal of the mold yielded raised conductive features crosslinked to an underlying layer.

### 2.3. Cell Culture

L929 murine fibroblasts and C2C12 murine myoblasts were cultured according to standard protocols in Dulbecco’s Modified Eagle Medium (DMEM) supplemented with 10 % fetal bovine serum (FBS) and 100 U/mL penicillin/streptomycin in an incubator at 37°C in 5% CO_2_. PF-AKR circular gels were soaked in sterile DI water to remove excess CaCl_2_ overnight. Gels were then soaked in 70% ethanol for 30 min, rinsed in PBS 3x and sterilized under UV light for 30 min. Cells were seeded at a concentration of 10,000 cells/well and given 24 h to attach. Media was changed every 2 days, and images were taken at Day 7. To assess viability, samples were removed on Day 7 and stained with a Live/Dead cell imaging kit (Thermo Fisher, R37601) for 15 minutes. Stained cells were imaged using a fluorescent microscope (EVOS FL Cell Imaging System, Thermo Fisher). Live and dead cell counts and percent viability were determined from the fluorescent images for each day using thresholding and particle analysis in ImageJ (at least n=5 frames captured at 10x magnification for each day).

Cells were seeded on micromolded hydrogels with patterned lines. After 10 days, samples were fixed in 4% paraformaldehyde (PFA). Fixed cells were prepared for fluorescent imaging using the following procedures, conducted at room temperature, followed by with 3 rinses with 1x PBS between each step. Cells were first permeabilized in 0.2% Triton-X for 10 minutes. Samples were then immersed in TRITC-phalloidin (Sigma Aldrich P1951, 50µg/mL) for 30 minutes to stain f-actin, followed by staining nuclei with for DAPI (Sigma Aldrich, D9542, 300nM) for 10 minutes. The cells were imaged (EVOS FL Cell Imaging System) and processed using ImageJ.

### 2.4. Electrochemical characterization of conducting hydrogel

Linear sweep voltammetry (LSV) was performed on conducting and non-conducing hydrogels (0, 0.25, and 0.5% w/v PEDOT:PSS, 3 replicates each) using a Gamry Interface 1010E potentiostat (Gamry Instruments, Warminster, PA) at a scan rate of 500 mV/sec over a potential window of −1V to +1V. Measurements were taken in triplicate immediately after gelation. To study the properties of films after diffusion of CaCl_2_, films were immersed in DI water to remove CaCl_2_. The water was replaced after the first 10 minutes, then every 30 minutes thereafter for two hours. Intermittent electrochemical readings could be used to assess the effect of trapped CaCl_2_, as the shape of the voltage vs current curve changed from curved to linear. Final readings of films with the CaCl_2_ removed were taken in triplicate for each sample. The average current vs. voltage curves for fresh and soaked conducting and nonconducting gels were plotted (n=9 for each curve, three samples x three readings each).

### 2.5. 3D Printing using aqueous PF solution

PF solution could be 3D printed via digital light processing utilizing a BIONOVA X 3D Printer system (CELLINK, Gothenburg, Sweden). To enable crosslinking at 405 nm, LAP photoinitiator (0.5% w/v in final solution) in DMSO (<50µL) and vortexed into a 7.5% w/v PF/AKR solution. 500 µL of the solution was pipetted into a well of the glass bottom (non-adhesive), black-sided 24-well BIONOVA X printing plate. The desired shape to be printed is loaded as an .stl file and sliced with the machine’s software. The slicing and printing parameters can be tuned depending on the target geometric and mechanical properties and printing material used. Here, print parameters were optimized for high fidelity 1 cm^3^ small footprint cubes and grids out of both conducting and nonconducting precursors (Layer height: 100 µm, light intensity: 60%, motion speed: 0.02 mm/s). The printing probe submerged in the photoactive material solution is used for direct in-well printing using UV light. Once cured, uncrosslinked solution was removed and transferred to the next well for subsequent prints up to three times, reducing wasted material. Multiple wells could also be filled with solution and different shapes could be printed in a programmed sequence. After removing the residual printing solution, the remaining crosslinked shapes were rinsed in DI water and easily removed from the well using tweezers. Depending on motion speed, the estimated time to print ranged from 1-5 minutes for cm-scale constructs.

## 3. Results

### 3.1. Formation of silk fibroin hydrogels

Photopolymerizable fibroin was first reported via addition of 2-isocyanatoethyl methacrylate to SF. With behavior similar to a negative tone photoresist, the methacrylated fibroin derivative -referred to as fibroin protein photoresist (FPP) or photofibroin (PF) – undergoes a transition from a liquid phase to a solid phase in the presence of crosslinking irradiation.[18] FPP/PF has similar secondary structure as SF, and was used to create water-insoluble micropatterned substrates.[31] Similarly, methacrylate/acrylated versions of silk fibroin (referred to as SilkMA, Sil-MA, FMA) hydrogels were synthesized by adding glycidyl methacrylate (GMA) to SF. PF’s higher degree of crosslinking makes it insoluble in water, with high stability in biological environments.[32] In prior work, the PF was dissolved in HFIP and FA as solvents which were ideal for forming extremely robust films and nano/microscale constructs via photolithography. The higher volatility of these solvents confer mechanical properties for flexible substrates which can support stress under deformation in a dynamic environment.[33] PF has applications in electrical and optical devices that take advantage of its precise, high-fidelity structures.[34] However, these solvents can be expensive and difficult to work with. Consequently, in this work, we endeavored to find a benign and water-based solvent for PF. Ajisawa’s reagent (AR) developed several decades prior, provides an appealing option as a water-based solvent for PF.[28] AR a ternary reagent is a mixture of CaCl_2_, EtOH, and H_2_O, with a molar ratio of 1:2:8 and has been successful at dissolving SF and PF rapidly. It is hypothesized that this is because the ethanol brings Ca^2+^ into the crystalline region.[35] Initially, the ability to form large scale hydrogels via soft lithography was shown (**Figure 2a**).

**Figure 2.**
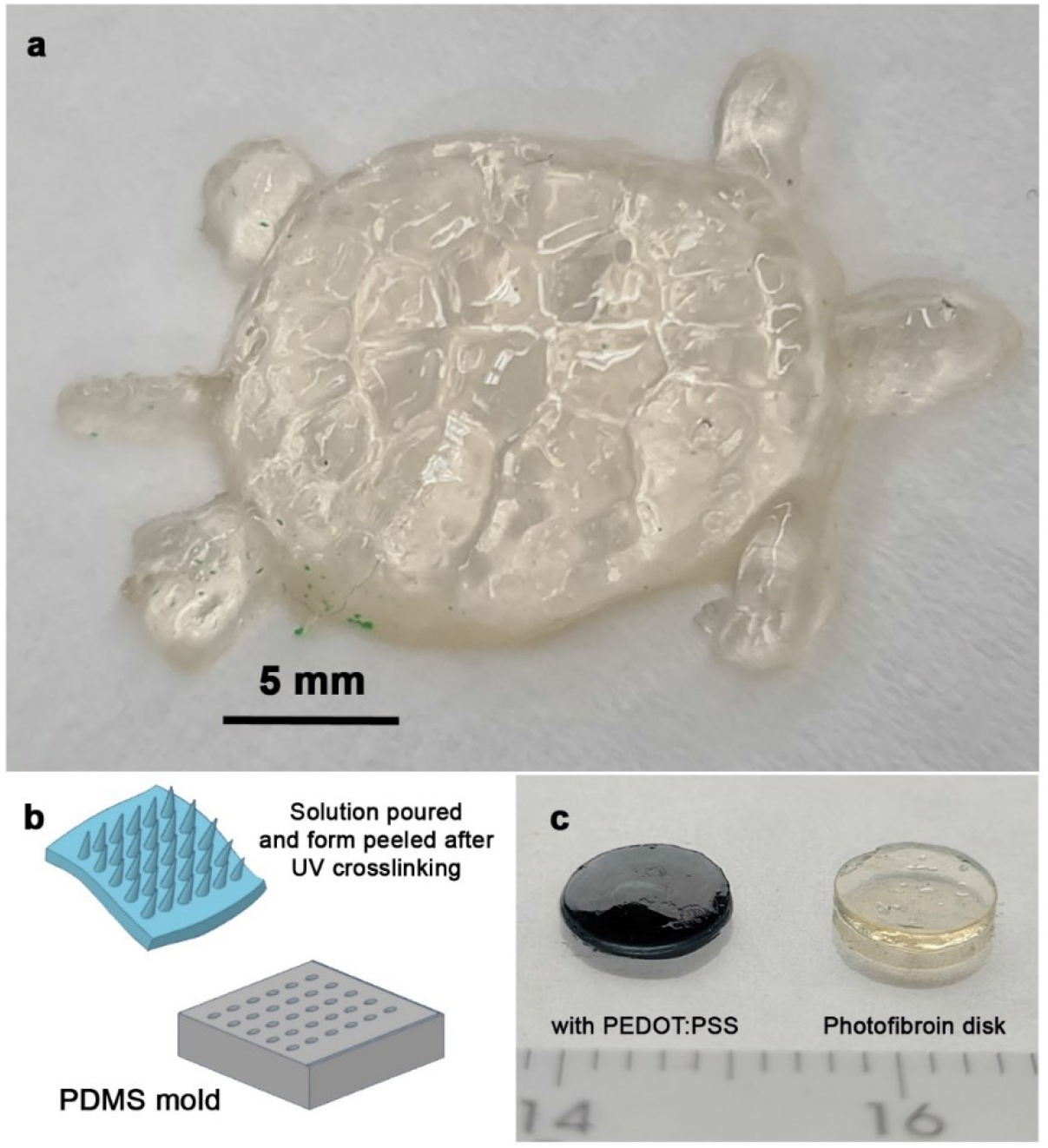
Formation of large-scale structures using soft lithography. The liquid precursor (photofibroin + photoinitiator) is poured into a PDMS mold with the negative design. Following UV crosslinking, the solid pattern can be peeled from the PDMS master (schematic shown in panel (b)). A large-scale turtle formed is shown in panel (a), with disks of the photofibroin with and without a conducting polymer (PEDOT:PSS) shown in panel (c).

Micromolding, or soft lithography is a technique where a hydrogel is cured in a negative mold to form designs on its surface. PF can be cured and removed easily from elastomeric PDMS micromolds, as well as commercial 3D-printable materials including ABS and photoactive resins while preserving microscale feature fidelity. The photoreactive polymer precursor solution was poured into PDMS molds and cured (crosslinked) by exposure to UV light (365 nm). The molds were easily peeled off to obtain the solid hydrogels (**Figure 2b**). The hydrogels were washed to reveal the high fidelity of the transferred features. Even complex hydrogels could be formed including conductive hydrogels (**Figure 2c**) and microneedles (**Figure 3a**). To form microneedle films, a two stage photopolymerization was employed in which thin films were first formed and placed over a mold with microneedles. Owing to the crosslinking between the two layers, an integrated, free-standing film with microneedles could be formed (**Figure 3a, b**). The films can be easily handled despite their soft nature (modulus on the range of 100s of kPa, data not shown).

**Figure 3.**
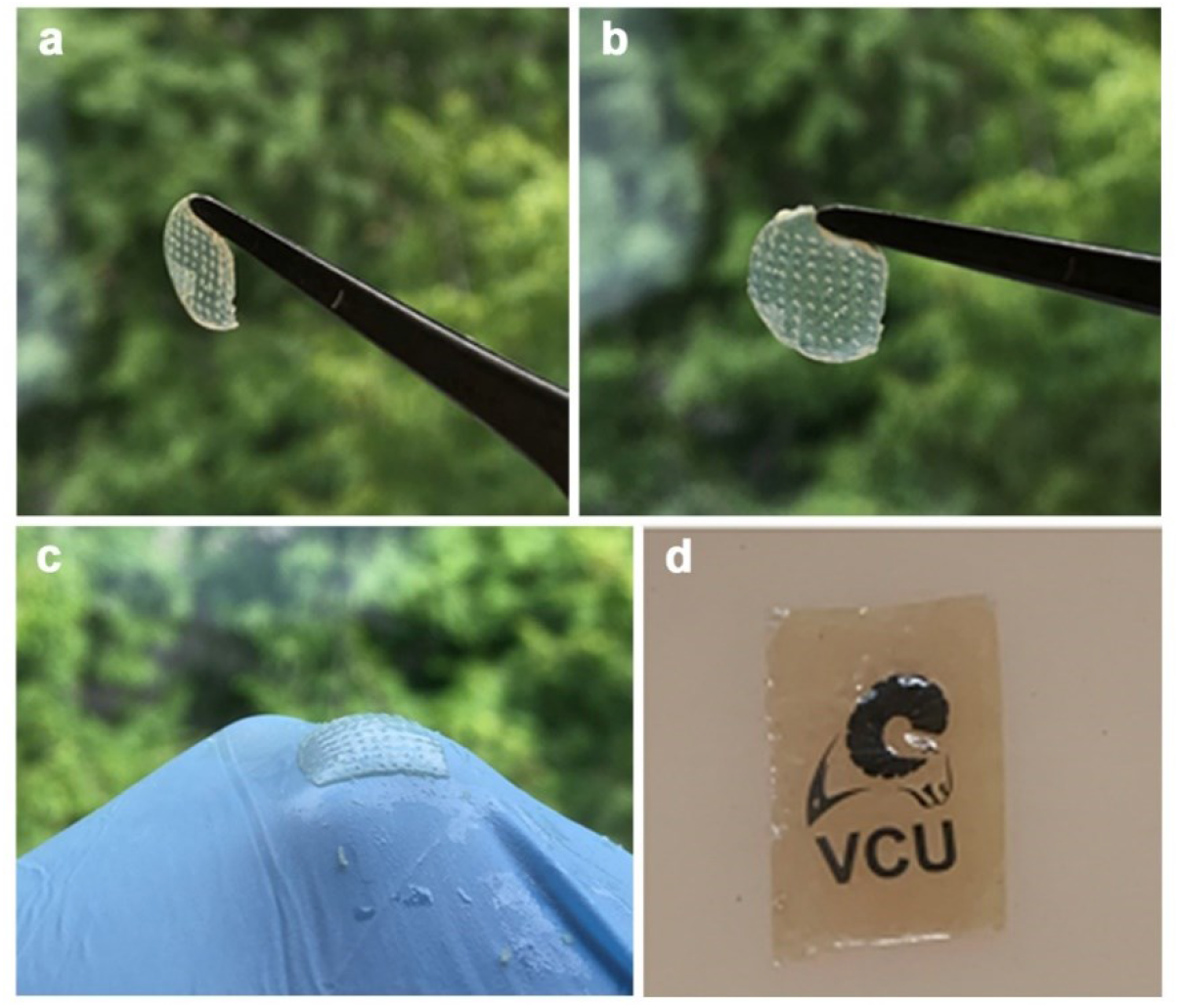
(a, b) microneedles formed using soft lithography (c) attachment to a glove is maintained under wet conditions. (d) Adhesive behavior of the ternary Ajisawa reagent whereby a printed photofibroin label is able to adhere to a synthetic skin model.

The hydrogel structures are extremely stable (for instance, the turtle shown in Figure 2 is stable in a buffered solution (PBS, pH 7.4) for more than 6 months without any loss of integrity) and can be mechanically tuned by controlling the degree of photopolymerization. The presence of the calcium further confers interesting mechanical and adhesive properties. Ca-modified silk obtained by dissolving SF and CaCl_2_ are known to be highly stretchable.[29–31] A transparent and stretchable Ca-modified SF was also demonstrated as a strong and biocompatible adhesive for epidermal electronics.[36] In the presence of Ca^2+^, the silk film becomes rich in random coils due to the metal chelation and water-capture, which results in a soft and pliable film. Similarly, the metal chelation provides cohesion and viscoelastic properties to the crosslinked film. High adhesion by large energy dissipation with good interfacial contact and cohesion enhances the peel strength. This is manifested by films becoming adhesive as shown by the ability to adhere to latex gloves (**Figure 3c**). Remarkably, the Ajisawa’s reagent can also be used as an adhesive (**Figure 3d**). In this demonstration, a thin film of crosslinked PF was imprinted with a bioink to form a readable label.[29] The use of the ternary solvent allows the silk label to adhere strongly to a synthetic skin model even under wet conditions.

### 3.2. Cell Culture on PF hydrogels and cytocompatibility

The biocompatibility, biodegradability, and low immune response of SF has been well demonstrated.[4, 31] Similarly, Sil-MA has found use in wound-healing and bioprinting hydrogels for encapsulating cells.[21] However, SF is known to lack cell adhesive domains, which often necessitates use of expensive supplements such as fibronectin to promote adhesion to fabricated constructs.[37] Due to the ability to form a wide diversity of architectures, PF represents a versatile biomaterial for tissue engineering and regenerative medicine.[6, 38] We had earlier shown the cytocompatibility of various forms of PF crosslinked from their HFIP and FA solutions.[31] Here, we explored whether the use of AR/AKR as a solvent contributes negatively in any manner to the previously established cytocompatibility.

The crosslinked hydrogel was explored as a cell substrate for two cell lines L929 murine fibroblast and C2C12 myoblasts. Cells were seeded and cultured over the course of 7 days. It was found that the PF/AKR hydrogels facilitate the robust adhesion and spreading of both cell lines (**Figure 4a, b**) . Cells exhibit flat morphologies and maintain high viability over the course of seven days, growing well on the hydrogel surfaces without the need for cell adhesive coatings or treatments. SF substrates formed using HFIP/FA, while non-cytotoxic, do not show a high level of cell adhesion and growth. In contrast, and most interestingly, the PF substrates with AR/AKR show excellent cell adhesion and growth (**Figure 4c)**. Our results were comparable to higher cellular adhesion on PF films that had been functionalized with polydopamine (PDA) (**Figure 4d**). It has been well established that PDA coatings improve cell viability, metabolic activity, and leads to flatter morphology, a function of increased surface roughness due to micro/nanoscale assemblies and modified organic surface chemistry. The ability of PF hydrogels formed using AR/AKR to replicate such adhesion with PF alone (no added adhesion promoter) shows that the solvent matters in terms of their use as excellent cell substrates. Live/dead staining of the cells indicated high viability, with >98% viability maintained across all days (**Figure 4e**). Thus these hydrogels are capable not only of providing excellent stability and mechanical properties, but also supporting cell viability and growth.

**Figure 4.**
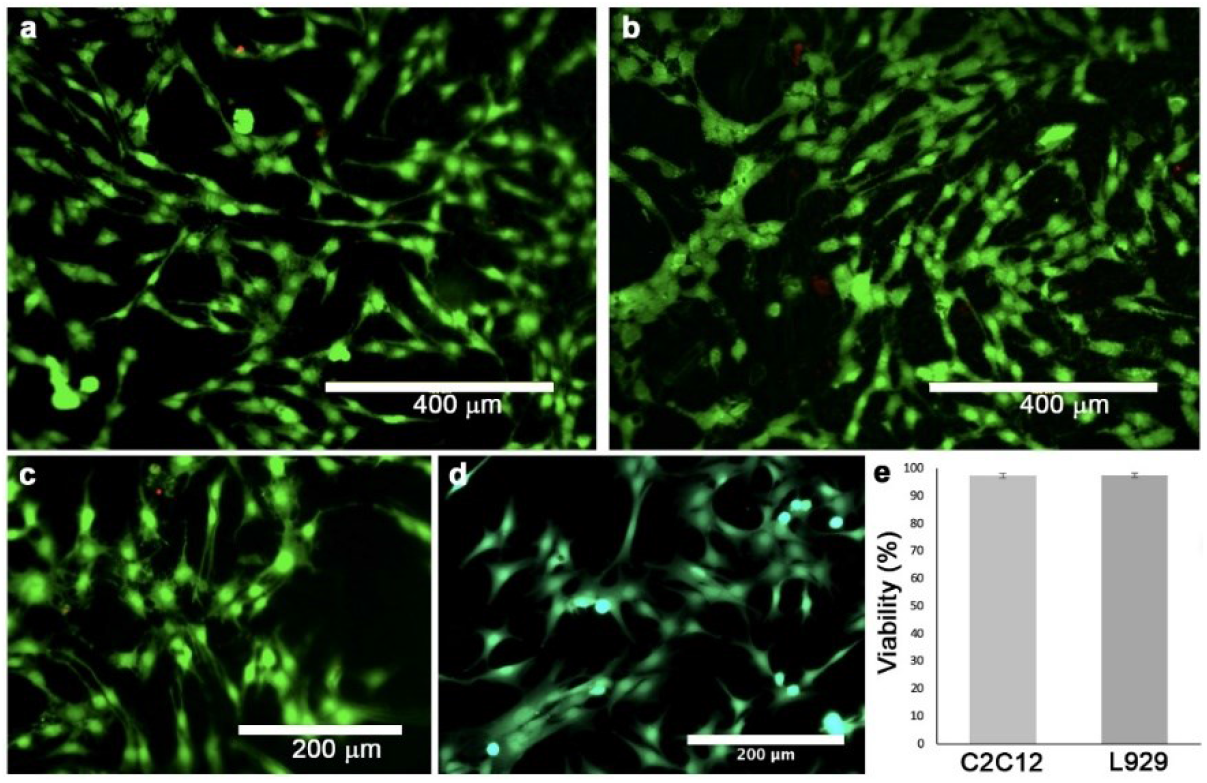
Cytocompatibility of PF gels on Day 7 for (a) C2C12 cells and (b) L929 cells (Scale bar = 400 µm). Cells proliferate to and spread over the hydrogel surfaces, maintaining high viability. (c) Close up of the C2C12 cells with a similar comparison to (d) PDA coated silk fibroin surface. The surfaces promote cell attachment and movement evident from long pseudopodia extensions with anchoring points on the surface (e) Quantitative measurements of cell viability and count over 1 week on the substrates. Error bars represent standard deviation with n=5 samples. Cells were stained using Live/Dead staining.

Soft lithography also provides utility to form hydrogel scaffolds with cell-instructive architecture, such as microchannels. Cells readily grow along patterned microscale grooves, with feature fidelity seen in parallel, clean gel edges (**Figure 5**). In fluorescent stained images, cell nuclei are visible spread evenly over the entire hydrogel surface, with f-actin cytoskeleton covering much of the area. Scaffolds not only support cell growth as they settle within grooves, but cells preferentially adhere upon the entire surface and spread over and along micropatterned features, indicating the ideal properties of this cell substrate. The easy patternability of PF can also be taken advantage to develop substrates with topographical cues for cell guidance and remodeling. 100µm lines were formed by micromolding. Two different cell lines - L929 fibroblasts (**Figure 5a**) and C2C12 myoblasts were seeded on these surfaces following the procedures as described above. In the case of cell types such as myoblasts, cellular alignment is an essential precursor to differentiation and development.[39] After 7 days, cell alignment within the troughs of the patterns was observed (**Figure 5b, c**). In the lower panel, C2C12 cells are shown at Day 0 and Day 7. Myoblasts appeared to align parallel to the micropatterns, elongating and appearing to fuse in patches. Fibroblasts also aligned along the patterns, and as a component of connective tissue, the precise patterning of fibroblasts could be useful in creating biomimetic constructs. Thus, micropatterned substrates can be explored in various configurations to influence cell arrangement, finding utility as cell-instructive substrates in tissue engineering.

**Figure 5.**
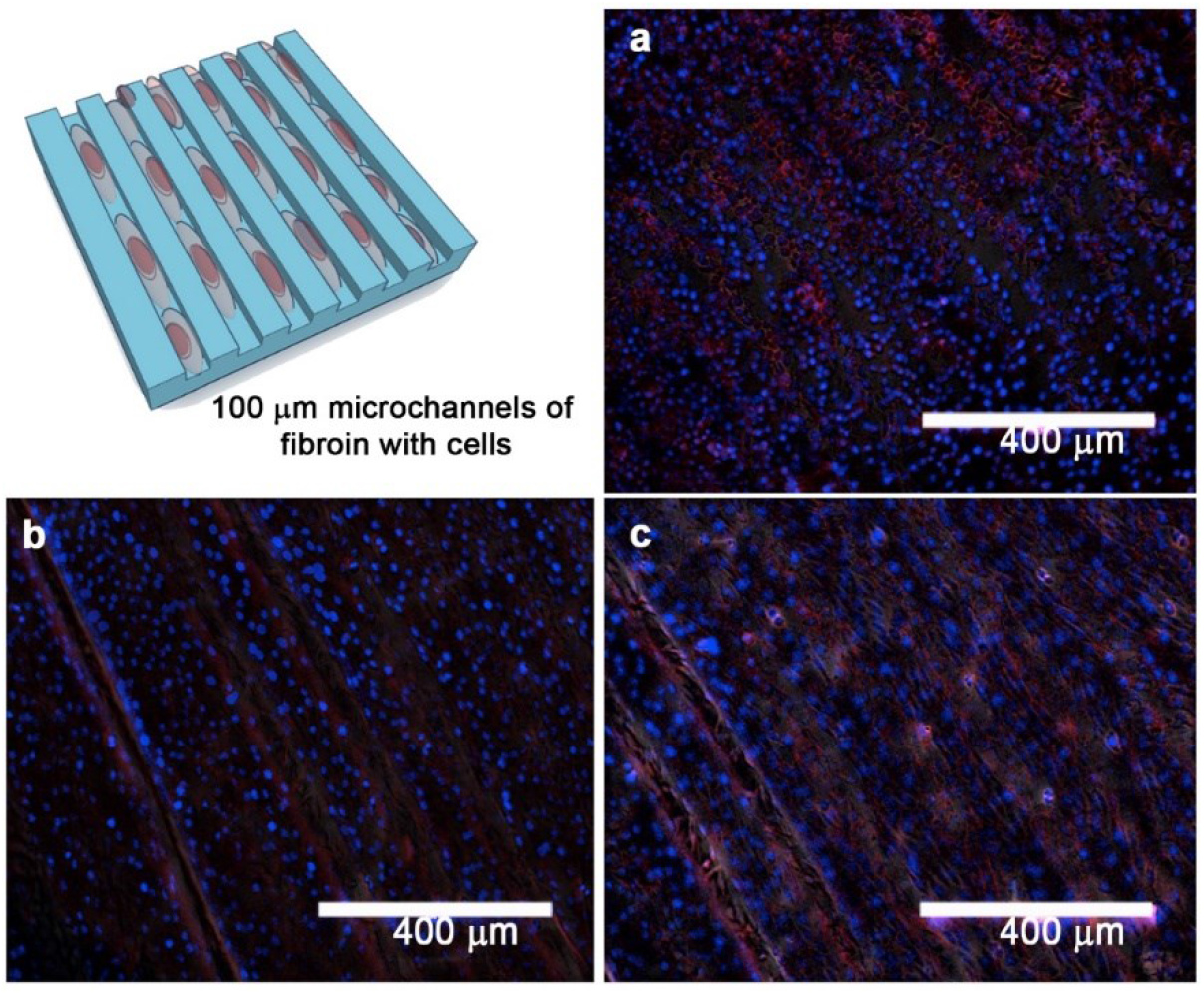
(a) C2C12 cell growth in the microfluidic channels fabricated using soft lithography. Cells (nuclei, blue) both settle within patterned grooves and grow to cover the microscale features, with f-actin microfilament network (actin, red) covering much of the hydrogel surface. For C2C12 cells, cells are aligned in the direction of the scaffold microarchitecture, and being to elongate and fuse along the channels. (b) L929 exhibit similar robust growth, evenly covering gels and aligning along patterned areas. Micromolded channels on gels exhibit precise patternability with high feature fidelity over mm to cm-scale areas.

### 3.3. Electrochemical Characterization

Soft electroconductive materials can provide avenues towards regenerative medicine as scaffolds that recapitulate aspects of the native cell environment and enable tissue-integrated bioelectronic sensing and stimulation. Few silk fibroin-based electrochemically active materials have been demonstrated, especially ones that are capable of being DLP-3D printed. Here, organic conducting polymer PEDOT:PSS could be dispersed in AR/AKR via ultrasonication in the amount of 0.25% or 0.5% w/v, in which PF is easily dissolvable. Prior work has introduced conducting elements into a regenerated silk fibroin matrix via immersion of a construct into a water/ethanol/conducting material co-solvent system, entrapping conducting material in the surface.[40, 41] One study was able to disperse PEDOT:PSS in CaCl_2_-formic acid with SF solution, with high Ca^2+^concentration maintaining random coil configuration and loose polymer networks, but the required use of organic acid for SF dissolution limits application.[42]

The conductivity of freshly made conducting gels (0, 0.25, and 0.5% w/v PEDOT:PSS, 3 films per condition) was analyzed using linear sweep voltammetry, then evaluated again after multiple wash steps in DI water. The gels have conductivity in the µA range, typical of conducting polymers. Increasing PEDOT:PSS concentration leads to an increase in conductivity across the tested range of voltage (**Figure 6a**). When films are freshly made, free Ca^2+^ ions work as current carriers within the polymer network and interact with themselves and the matrix to form complexes, resulting in nonlinear behavior and higher currents across all formulations. Since most applications involve aqueous environments, evaluating the matrix after soaking and washing in DI water was evaluated. After two hours of successive water changes, gels exhibited a linear current response to increasing voltage, demonstrating stability of concentration gradients and matrix interactions, with slightly lower overall conductivity a result of free Ca ions being removed. Incorporating larger amounts of conducting PEDOT:PSS increases the conductivity of the hydrogels beyond the amount seen due to presence of water and Ca ions in the nonconducting case. The discernable changes in material conductivity seen in this hydrogel material show practicality for biological electronic applications in consumer devices or clinical products, and demonstrate the novel ability to create bulk conducting composites of PF. Conducting circuits (PF+PEDOT:PSS) could be formed on flat PF substrates in a two-step process resulting in integrated biodevices. The circuit was cast on a pre-formed thin film of PF. Photopolymerization resulted in fusing of these two layers to form a unified construct as seen in **Figure 6b**.

**Figure 6.**
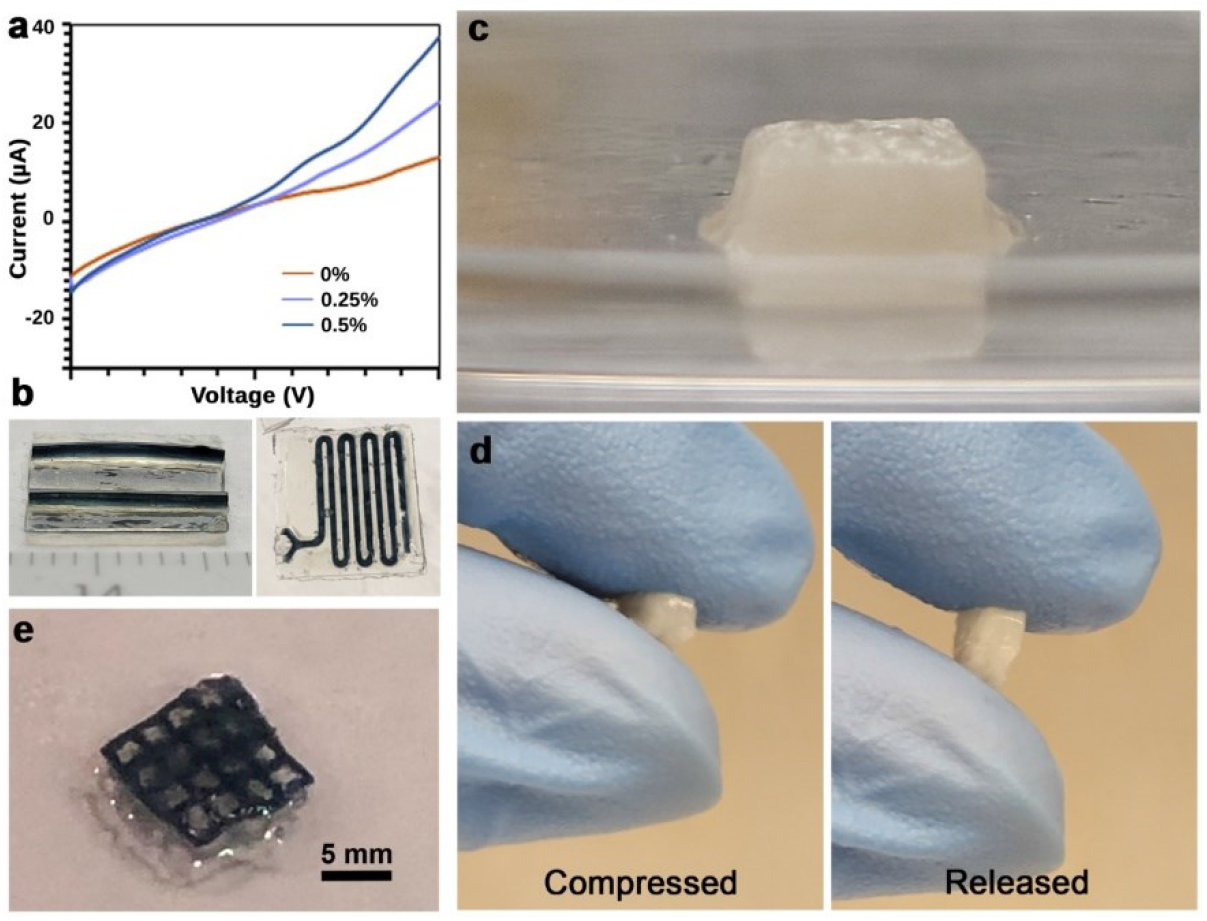
(a) Electrochemical characterization of the conducting hydrogels using linear sweep voltammetry (different % compositions of PEDOT:PSS in the hydrogels are indicated). Hydrogels were soaked in water prior to measurement. (b) Conducting patterns (PF+PEDOT:PSS) formed on flat PF substrates (c) 3D printed hydrogel of 1 cm x 1 cm x 1 cm cube with internal voids via DLP (d) 3D printed structures can be compressed and released to regain their original shape (e) 3D printed conducting hydrogel pattern comprising PF and PEDOT:PSS.

### 3.4. DLP 3D printing of photofibroin hydrogel

SF is difficult to directly 3D print due to its low viscosity, and previous methods have utilized methanol-induced crosslinking or forms of suspension printing to provide mechanical support.[43] As opposed to extrusion, low material viscosity is desired in vat polymerization. SF composites incorporating PEG-DA and other photocurable materials have been utilized in DLP printing, at the cost of reduced flexibility, and potentially, biodegradability.[44-46] Thus, photofibroin with its ability to be photopolymerized is uniquely positioned as a biomaterial for DLP printing.[47] Interestingly various components balance protein solubility and aggregation in the silk native gland, with Ca^2+^ ions shown to prevent protein alignment and aggregation in unspun silk.[48] Previously, PF in HFIP/FA with remarkable nanoscale fidelity and high mechanical strength was limited by compatibility with equipment, safety and sustainability in the large amounts needed in vat polymerization.[7, 49] The CaCl_2_-containing AR/AKR solvent used in this study is low-viscosity, and facilitates homogeneous dissolution of PF at amounts exceeding 10 w/v%.

The BIONOVA X DLP 3D printer system was utilized in this study. A 3D printing probe directly cures the desired construct within a small volume of precursor solution within the wells of a standard well plate. This set up is easier to operate in comparison to traditional DLP or stereolithography plates that move within a large vat of photoresin. 7.5 w/v% solutions of PF in AKR, with 0.5 w/v% LAP photoinitiator enable printing at 405 nm light used with this printer. 500 µL of printing solution was pipetted into wells of a 24-well plate, and the slicing and print parameters were varied to find an optimal setting for test shape formation (Layer height: 100µm, Light intensity: 60%, Motion speed: 0.02mm/s). A 1 cm^3^ with 100 µm sized features (internal mesh grid) was printed (**Figure 6c**) within 1 minute - a relevant time frame for rapid prototyping and bioprinting with encapsulated cells. The 3D printed structures were washed with DI water to remove unreacted PF, LAP and CaCl_2_ and could easily be removed and manipulated with tweezers. Our 3D printed silk hydrogels with microscale internal architecture are robust and can be handled without any special tools (**Figure 6d**). As shown in the figure, it exhibits recoverable compressive ability, and can be depressed with a finger or spatula, returning to its original height. A video showing this compression and release is shown in the supporting information as **Movie S1**. Gels can be wetted and dried with minimal swelling, with defined cube edges and height uniformity. It is important to note that various parameters such as stiffness can be further fine-tuned depending on desired properties for different applications, including as cell scaffolds or structural devices. In comparison, it is worth noting that Sil-MA formed from fibroin methacrylation by glycidyl methacrylate used in DLP printing, increased curing time, photoinitiator and Sil-MA content resulting in stiffer gels due to a more stable hydrophobic domain.[21] Print parameters can be iteratively tuned to balance efficiency versus feature resolution.

The conducting PF could also be printed as conducting grids (**Figure 6e**). To the best of our knowledge, the DLP printing of conducting organic polymers such as PEDOT:PSS within the silk matrix has not been demonstrated before, due to solvent incompatibility and aggregation. At 0.25 w/v% PEDOT:PSS dispersed in the solution, an increased viscosity and marked decrease in light transmittance is noted.[50] However, conducting lines of 100 µm thickness are still easily formed. The ability to spatially pattern microscale electronic features is of much interest for applications in biosensing, stimulation, and cell culture. The changes in optical and rheological properties of the conducting bioink provide future opportunities for tuning of optimal print parameters.

## 4. Discussion

Silk fibroin (SF) and its composites have long been studied for their use in tissue engineering, drug delivery, and regenerative medicine.[5] Despite a versatile palette of properties, enhancements to SF have typically required composites with both synthetic and natural polymers including PVA, polyurethanes, chitosan, gelatin etc.[15, 16] Chemical cross-linking methods provide benefits for SF hydrogel fabrication such as reproducible gelation, network structure control, and mechanical properties. Limitations including solution stability and spatial and temporal control still remain.[51] Alternatives such as light are sought - in particular, the photocrosslinkable variants of silk fibroin (known variously as photofibroin (PF), fibroin methacrylate (Sil-MA, SFMA) etc.) induce precise spatial and temporal control of properties. As noted above, our group and others have developed such biomaterials for applications including tissue engineering (as high-resolution 3D scaffolds to provide a supportive environment for cell growth and tissue regeneration), hydrogels (wound healing, drug delivery and matrices for cell encapsulation), customizable biomedical devices, microfluidic devices, and biosensors, and bioadhesives and sealants.

### Synthesis of PF using IEM

While a number of different synthesis strategies have been devised to date, following our initial work using 2-isocyanatoethyl methacrylate (IEM) to form photofibroin (PF),[18] there are some distinct advantages to this form of photocrosslinkable fibroin (PF). Since IEM is an electrophilic compound, it can react with both weak hydroxyl groups and strong nucleophiles (amino groups in lysine and arginine residues) at comparable rates. On the flipside, unlike the MA reaction, which can be conducted in an aqueous medium, methacrylation by IEM is carried out in anhydrous dimethyl sulfoxide (DMSO). Both GMA and MA utilize the 0.2 mol% lysine residues for introducing methacryloyl groups into the SF chain. GMA can attach two methacryloyls to one amino group, leading to a more effective reaction in comparison to MA.[24] Due to the capability of reactive IEM, higher degrees of methacrylation are obtained than in the case of MA, with values up to 9.3% reported. Thus, the PF formed via IEM is extremely versatile at forming structures of extraordinary fidelity and strength. Sil-MA hydrogels tend to be weaker resulting in lower resolution and strength particularly in multilayer or large volume constructs.[25] The PF product formed is easily lyophilized into a powder, highly shelf-stable for several months and is extremely reproducible from a variety of different cocoons. However, to date, PF use has been limited to forming solutions using (hexafluoroisopropanol) HFIP or formic acid (FA) (**Figure 1**). These have been excellent at *photolithographic* routes of fabrication and the solvents do not affect the bio/cytocompatibility of the constructs. However, the use of volatile and expensive HFIP/FA can also be a deterrent to wider use. The utilization of PF via the sustainable and low cost ternary aqueous solvent of water-ethanol-CaCl_2_ can open new avenues for its use.

### Improvement in mechanical properties due to calcium

SF is exceptionally strong, but tends to be brittle when dry, limiting its applications. The presence of CaCl_2_ in the ternary mixture of Ajisawa reagent provides an added benefit to the mechanical properties of the biomaterial. Within the silkworm itself, Ca^2+^ ions impede silk molecular alignment and aggregation, mimicking the natural storage conditions of unspun silk in the silk gland.[48] CaCl_2_ has been shown to decrease Young’s modulus and increasing elongation of SF at break.[52] While Ca ions have a negligible effect on hydrogen bonds in the silk matrix and secondary structure, the increased water uptake decrease crystalline strength, while introducing structures conducive to stretchability.[53] The CaCl_2_ increases hydrophilicity owing to the Ca^2+^ ions which interact with the coil chains of silk as cross-linkers by chelation effect increasing water capture.[36, 54] Further plasticization via addition of CaCl_2_ significantly improves water content of films in environments containing the same relative humidity.[53] Overall, the hydrogels are extremely robust and mechanically strong (modulus on the order of MPa). They resist deformation and can easily return back to their original shape following compression as shown in **Figure 6d** and **Movie S1**.

### Cytocompatibility and enhanced cell adhesion

The cytocompatibility of PF substrates formed using HFIP and FA was previously shown. Therefore, the cytocompatibility of PF hydrogels formed using AR/AKR is an expected result. The most interesting take away from these experiments was the increased cell adhesion to the PF when using the ternary water-ethanol-CaCl_2_ solvent. To date, various cell types have been grown on PF films, including C2C12, HDFs, N2A, L929, and hBM-MSCs.[17, 31, 55] However, without pretreatment in media or fibronectin coating (expensive), cells frequently have low proliferation and nonspecific adhesion. SF heavy chains and crystalline β-sheets are hydrophobic, deterring cell attachment and reducing proliferation rates.[56, 57] Based on initial studies of cell attachment on PF films, fibronectin was needed to promote cell attachment. As shown in Figure 4d, the use of polydopamine as an adhesion promoter was also shown. Here, we hypothesize that the soft, porous, 3D matrix of PF hydrogels better facilitates cell interaction and integration between cells and the gel matrix in comparison to the more crosslinked films formed using HFIP/FA.[58] SF treated with CaCl_2_ has been shown to have rougher surface topography and improved hydrophilicity, which improves cell anchoring.[59] For the PF in AR/AKR, winding ridges are formed on the surface due to initial film formation and tension during the UV-curing process as the rest of the gel cures, providing topography for cells to settle and grow along. Patches of cells can be seen initially aligning along curving paths, then easily interacting over these features with the rest of the cells on the surface. Additional nano/microscale roughness is evident from cell pseudopodia extensions and anchoring at visible points on the surface. In addition to physical effects, the incorporation of CaCl_2_ and ethanol with SF has unique synergistic effects for cell culture. Calcium ions are essential to a range of native cell functions and communication, and an optimal low concentration has been shown to improve proliferation in engineered cell matrices that recapitulates the amount maintained in the cytoplasm. [60] For SF, a previous study confirmed that even after extensive dialysis (> 7 days), there is still a small amount of CaCl_2_ residue (about 1000 mg/mL) bound to SF treated with AR, relevant to cell culture where constructs are washed, sterilized, and immersed in aqueous media for days to weeks. [59] These residual calcium ions can both improve cell membrane stabilization, and alter the material properties of SF including altered electronegativity compared to materials prepared in HFIP or FA, which improves protein adsorption and cell binding, and amino acid interactions that influence secondary structure. Ethanol further allows the movement of Ca^2+^ into the beta crystalline regions of the fibroin, and formation of a more ordered crystalline structure. [61] Low concentration ethanol treatment have also been shown to promote a swollen hydration layer on the outermost micrometer-scale surface of silk fibroin films, improving cell adhesion and proliferation.[62]

### 3D printing of PF

Aqueous systems are more environmentally and biologically friendly and needed for compatibility with many types of fabrication processing equipment. The robust nature of the resulting gels enables compatibility with biofabrication techniques for more useful and representative structures. The use of PF with soft lithography can be used to form a wide range of features including large scale objects, microneedle patches, microfluidics, and electrical circuits. The ability to manipulate these micropatterned constructs without fragility is critical for clinical and commercial product viability, as transdermal patches, or subcutaneous implants. The true enhancement of the biofabrication comes from the use of SF in 3D printing.[63] As a printing material, it offers better mechanical strength than other natural biomaterials, while retaining benefits of controlled biodegradation and cell viability. However, owing to its low viscosity (46 mPa·s at 16% (w/v), 71 mPa·s at 24% (w/v)), it has not been useful for nozzle-based printing.[47]

These methods are limited by material rheological properties, and have limited ability to form complex 3D structures with voids and overhangs depending on self-supporting ability and adhesion between print layers. Nozzle-based printing also takes longer due to material filament extruded to build each construct, and nozzle size limits resolution. Alternatively, light-based methods such as digital light printing (DLP) use a projected UV light source to cure entire layers at once within a vat of (even low viscosity) photopolymerizable material. Thus, printing speed is increased and only depends on the height of all constructs, and enables complex geometries in high resolution. The use of light together with the photocrosslinkable SF opens new avenues in throughput, microscale resolution, complex spatial geometry, and curing kinetics. As shown above, the use of PF in the AR/AKR solvent can be combined with novel 3D printers that can utilize a small volume (e.g., microwell plates) or parallelized biofabrication approaches that can be directly combined with cell culture.

### Conducting PF hydrogels

While bulk aqueous dispersion of PEDOT:PSS was earlier shown with water-soluble photosericin,[29] attempts to incorporate conducting polymers with photopolymerizable silk fibroin in an aqueous system have been limited.[19] The use of conducting polymers provide higher mechanical and electrochemical stability in comparison to other additives to form electroconductive bioinks (e.g. Sil-MA and reduced graphene oxide (rGO)).[21] The aqueous conducting PF/AKR solutions thus enable us to combine the benefits of electrochemical activity with the facile micro/nanoscale light-actuated processing and mechanical strengths conferred by silk photofibroin. The prepared gel precursor solutions are more viscous than nonconducting versions with the same solid content, as calcium ions facilitate crosslinking of PSS via electrostatic interaction. However, the precursor solution is still homogeneous and easy to handle (e.g. 0.25% PEDOT:PSS solution used in this study).[64] The solution crosslinks in the same time frame under UV light (seconds), yielding gels with increasing dark blue color with increasing PEDOT:PSS concentration. As shown in **Figure 6a**, the electrochemical properties of the conducting hydrogels are very competitive in comparison to prior studies as well as temporally stable even in aqueous solutions.

## 5. Conclusions

In summary, this study demonstrates the successful development of photocrosslinkable silk fibroin (PF) gels using eco-friendly solvents, expanding the potential of silk fibroin for advanced biomedical and bioelectronic applications. By utilizing Ajisawa’s reagent as a sustainable solvent, this work addresses the limitations of traditional crosslinking methods, providing a safe and versatile alternative for silk fibroin processing. The photocrosslinkable nature of PF allows for precise control over material properties, enabling the fabrication of complex structures through light-based techniques such as 3D printing and soft lithography. While Sil-MA has several useful advantages, the PF shown has high mechanical strength, easy reproducibility and higher long term stability. The integration of conductive polymers like PEDOT-PSS further enhances the applicability of PF gels in bioelectronics, while the demonstrated cytocompatibility supports their use in tissue engineering and regenerative medicine. Overall, this research represents an enhancement of PF properties towards sustainable, high-performance biomaterials that are both biocompatible and adaptable to a wide range of medical and technological applications. Future work may focus on optimizing mechanical properties and exploring additional biofabrication techniques towards complex constructs for biomedical applications.

