## Supplementary figures and images for "Enhancing the versatility of photocrosslinkable silk fibroin using an eco-friendly solvent"

### Supplementary Movie S1

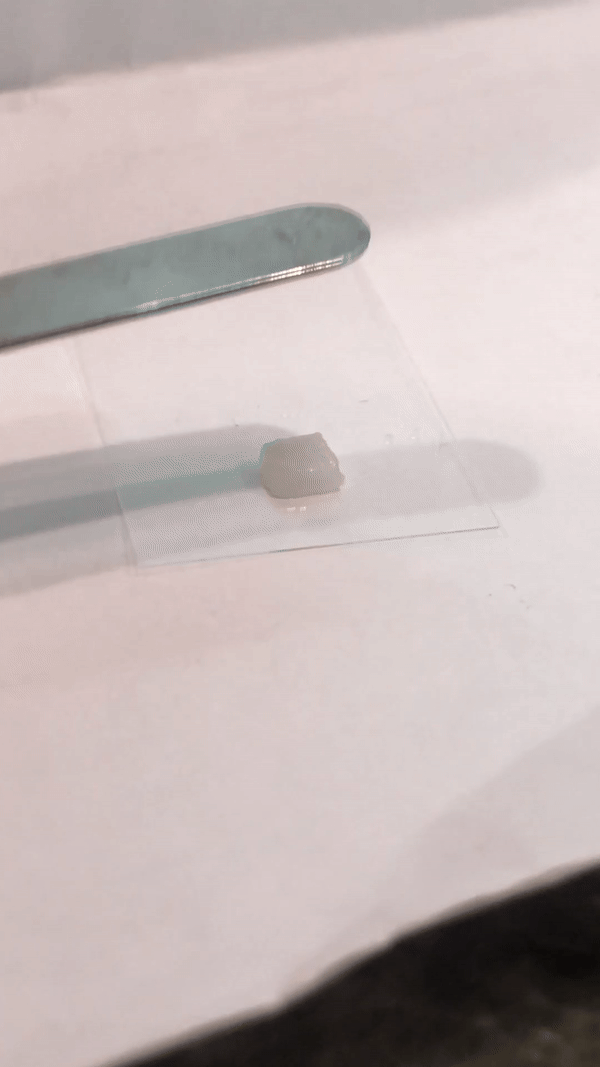
